# BoMBR: An Annotated Bone Marrow Biopsy Dataset for Segmentation of Reticulin Fibers

**DOI:** 10.1101/2024.10.02.616389

**Authors:** Panav Raina, Satyender Dharamdasani, Dheeraj Chinnam, Praveen Sharma, Sukrit Gupta

## Abstract

Bone marrow reticulin fibrosis is associated with varied benign as well as malignant hematological conditions. The assessment of reticulin fibrosis is important in the diagnosis, prognostication and management of such disorders. The current methods for quantification of reticulin fibrosis are inefficient and prone to errors. Therefore, there is a need for automated tools for accurate and consistent quantification of reticulin. However, the lack of standardized datasets has hindered the development of such tools. In this study, we present a comprehensive dataset that comprises of 201 **Bo**ne **M**arrow **B**iopsy images for **R**eticulin (BoMBR) quantification. These images were meticulously annotated for semantic segmentation, with the focus on performing reticulin fiber quantification. This annotation was done by two trained hematopathologists who were aided by Deep Learning (DL) models and image processing techniques that generated a rough automated annotation for them to start with. This ensured precise delineation of the reticulin fibers alongside other cellular components such as bony trabeculae, fat, and cells. This is the first publicly available dataset in this domain with the aim to catalyze advancements the development of computational models for improved reticulin quantification. Further, we show that our annotated dataset can be used to train a DL model for a multi-class semantic segmentation task for robust reticulin fiber detection task (Mean Dice score: 0.92). We use these model outputs for the Marrow Fibrosis (MF) grade detection and obtained a Mean Weighted Average F1 score of 0.656 with our trained model. Our code for preprocessing the dataset is available at https://github.com/AI-in-Medicine-IIT-Ropar/BoMBR_dataset_preprocessing.

## 1. Background

Reticulin fibers *are* an integral component of the bone marrow’s extracellular matrix, providing essential support for hematopoiesis, the process of blood cell formation. These fibers are visualized using silver impregnation techniques, revealing a delicate network of thin, uniform strands. The organized reticulin meshwork serves as a structural framework within the bone marrow environment. However, in several pathological conditions (Norén-Nyström et al., 2008; Fu et al., 2014; Buesche et al., 2008), there is an increase in reticulin deposition within the bone marrow, contributing to myelofibrosis. Separating and decoding the structural complexity of reticulin fibers, which are stained and examined under a microscope, from the Bone Marrow Trephine (BMT) images is labor-intensive and time-consuming (for an example of a BMT image and the corresponding complex reticulin fiber structure refer to Fig. 1). This process is also subjective and prone to the presence of inter-observer variability (Teman et al., 2010) and can, therefore, differ across pathologists (Lucero et al., 2016). Accurate grading of reticulin fibrosis is crucial and inconsistencies impact the accurate diagnosis, prognostication and treatment decisions.

**Figure 1.**
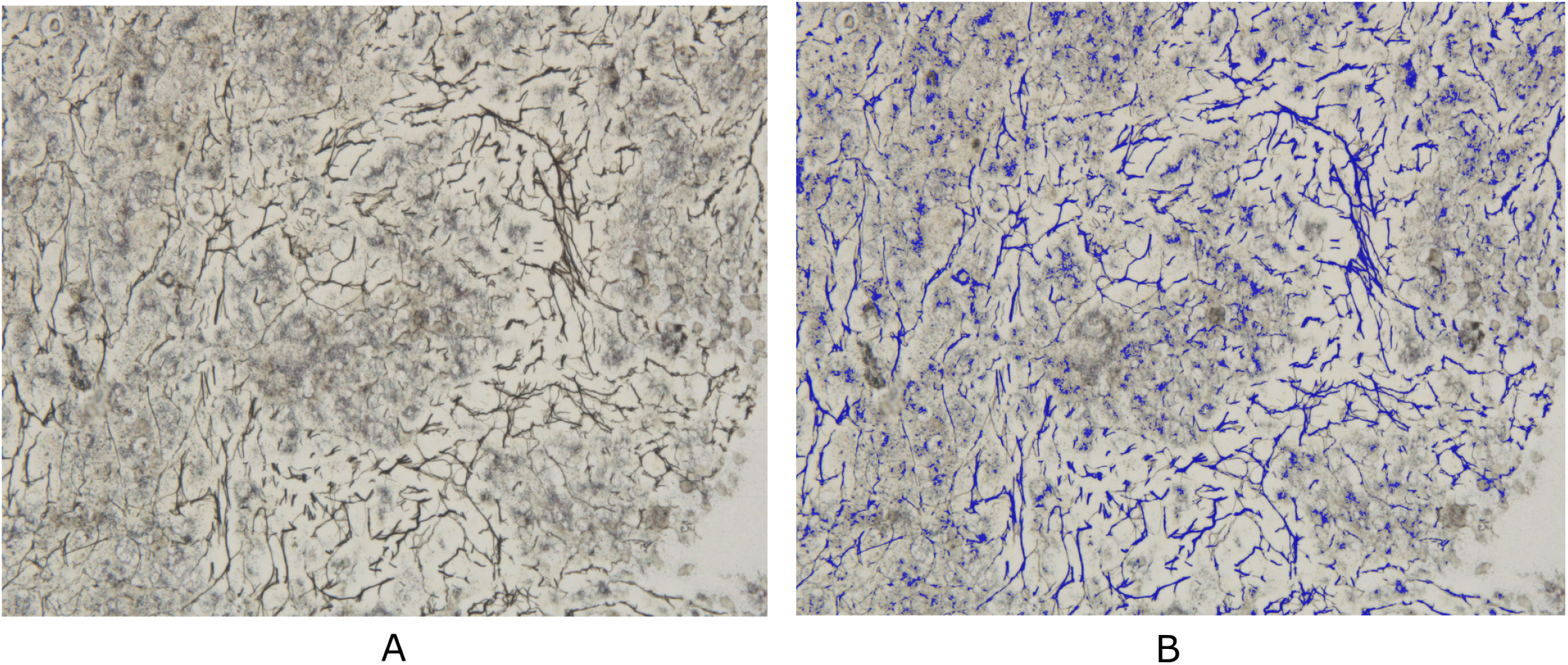
Sample of a BMT image from our dataset. A: Original unsegmented image B: Annotated image with reticulin fibers labeled in blue color

**Figure 2.**
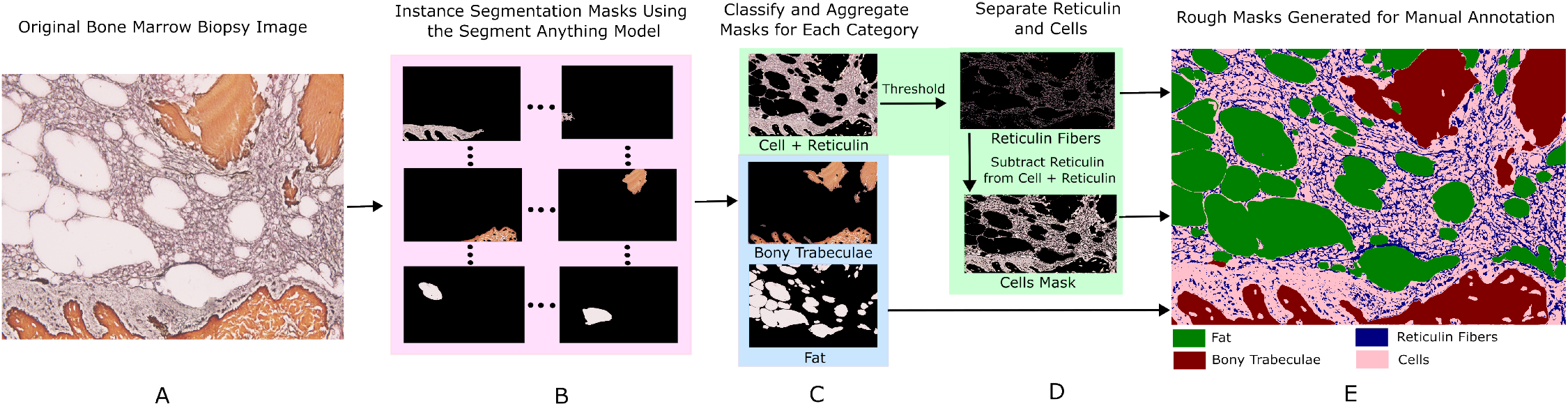
Flowchart representation of automatic annotations done by SAM. A: Original Image. B: Generating masks and classifying them: Masks with pixel intensity in the orange color range are classified as bone masks, masks having consistency in color among the remaining masks are considered as fat masks, and the remaining masks are categorized as cell masks. C: Masks of every category are added together to form one mask per category. D: Performing thresholding and shape filtering to isolate reticulin, resulting in a reticulin mask and a cell mask without reticulin. E: Annotations generated using SAM and classification.

**Figure 3.**
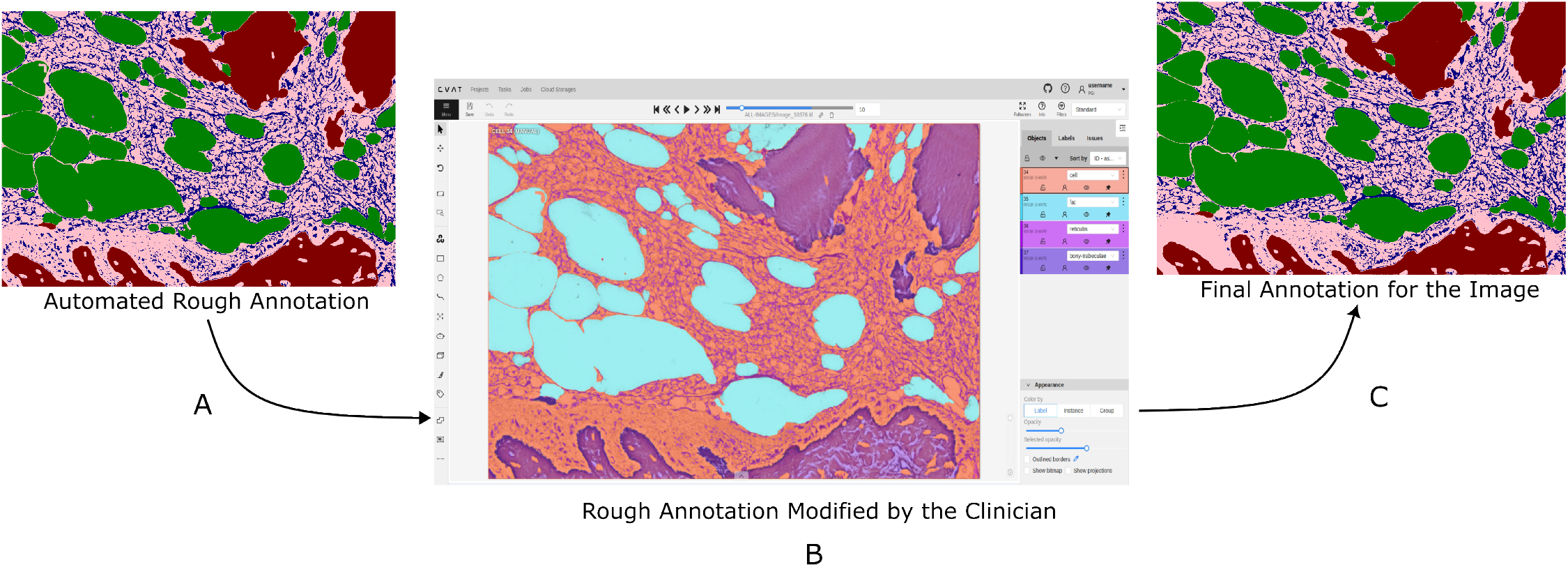
Procedure for annotation by experts. A: Automatic annotation using SAM and classification of masks. B: The pathologist uses Computer Vision Annotation Tool (CVAT) for refining annotations. C: Final annotation

In the past two decades, the advancements in obtaining digital images in the pathology coupled with the developments in deep neural network models for computer vision related tasks have led to multiple tasks in pathology being automated (for a review, please see (Amerikanos and Maglogiannis, 2022)). Consequently, there have been previous attempts for developing computational models for reticulin detection from BMT images. However, reticulin fibers exhibit variations in color intensity and distribution, making accurate segmentation and quantification difficult (Lucero et al., 2016). For example, Teman et al. (2010) developed a computer-aided solution that utilizes color deconvolution techniques to isolate the reticulin fibers from the image and then manually grade the MF. This work, however, ignores the complex polychromatic nature of reticulin fibers and is not fully automated thereby having a manual grading component.Lucero et al. (2016) quantified bone MF in a mouse model of myelofibrosis. However, its applicability to human bone marrow biopsies and more complex disease presentations requires further validation and adaptation. More recently, Ryou et al. (2023) introduced Continuous Indexing of Fibrosis (CIF), a DL approach aimed at enhancing the quantification and monitoring of fibrosis in using bone marrow samples. CIF utilizes a continuous 0-1 scale, departing from the traditional discrete 0-3 grading system, thereby improving fibrosis assessment in Myeloproliferative Neoplasms (MPN). However, a potential drawback is that this new grading scale may not align with established clinical practices, posing challenges for clinicians in interpreting and applying the results in standard clinical settings. As a result, these computer-aided approaches may not provide reliable and reproducible results, further highlighting the need for alternative methods that can overcome these limitations and offer more robust assessments of bone MF.

It should be mentioned that we were unable to locate any publicly accessible annotated datasets for the quantification of reticulin in BMT pictures. Ryou et al. (2023) made use of a private dataset that wasn’t accessible to the general public.

## 2. Summary

We propose the **B**one **M**arrow **B**iopsy images for **R**eticulin (BoMBR) dataset containing 201 BMT pixel-wise annotated images. Besides the pixel-wise annotation masks, we give information regarding the grade of MF, percentage cellular area covered by reticulin, and the average value of the elongation factor of the reticulin fibers in the image. The percentage of cellular area covered by reticulin indicates differences in the amount of area covered by reticulin fibers across different grades, while the elongation factor helps understand changes in the shape of reticulin fibers as the grade increases. This is by far the first such publicly available resource for annotated BMT images that focuses on reticulin fibrosis both globally and in the context of the Indian subcontinent. Our dataset aims to facilitate the effective and objective quantitative measurement of reticulin fibers, moving beyond the current qualitative, observer-dependent scoring systems. Beyond its primary application in the grading of myelofibrosis, the dataset may also serve as a benchmark for further studies. This includes its potential to assist hematopathologists in the classification of MPN, assessment of disease progression, and quantitative monitoring of the effects of myelofibrosis reversal in patients, particularly in the context of clinical trials and novel targeted therapies.

Further, to show the utility of the dataset, we train a DL model and show that the BoMBR dataset can help in training precise segmentation models that can further help in determing the MF grade of the BMT biopsy images. The BomBR dataset is preprocessed, split into train and test sets (stratified for the MF grade) and ready for use BoMBR: Bone Marrow Biopsy Dataset for Segmentation of Reticulin Fibers by researchers for computational processing. The code for designing the corresponding DL models is available on https://github.com/AI-in-Medicine-IIT-Ropar/BoMBR_based_models/tree/main.

The following sections of this paper provide a comprehensive overview of our work. We begin by discussing the significance and limitations of the presented dataset (section 3). Next, we highlight potential applications and license related details of the dataset in machine learning and clinical imaging (section 4). Following this, we describe the methodologies used for data collection (section 5.1) and the detailed annotation process (section 5.2). Finally, we demonstrate the use of the BoMBR dataset for predicting fibrosis grades, showcasing the potential of our dataset in aiding clinical diagnosis and research (section 6).

## 3. Discussion

The annotated dataset of BMT biopsy images presented in this study provides a valuable resource for quantitative assessment of fibrosis severity in hematological disorders. By leveraging automated segmentation followed by expert annotation, we have created a comprehensive dataset covering a range of MF grades (MF-0 to MF-3). This dataset includes detailed annotations for reticulin fibers, bony trabeculae, fat regions, and cellular region, facilitating in-depth analysis of bone marrow pathology.

Despite the strengths of this dataset, there are some limitations to consider. Firstly, although the annotated dataset will aid in the quantification of reticulin fibers, it does not identify the presence of collagen fibrosis and osteomyelosclerosis, which are important components of the overall fibrous matrix. Secondly, the dataset has not been prospectively validated for use in clinical laboratory settings, including its applicability to whole slide images (WSI). Additionally, how clinicians and hematopathologists should interpret the resultant data has not been studied.

Additionally, further research is being conducted to test these models using Whole Slide Images in clinical situations. To verify the usefulness and efficacy of these models, prospective studies should evaluate how well they function in actual diagnostic workflows. Multi-centric setups should be explored to include more images and train more generalizable models. In future, the dataset can be expanded to enable the prediction of specific types of MPN and not just the MF grade. In order to perform annotation for situations where the trained models underperform, semi-supervised setups for MF grade prediction should be investigated on the DL side.

In conclusion, the annotated dataset of BMT biopsy images represents a significant advancement in the field of hematopathology and automated diagnostic tools. It provides a foundation for developing robust DL models capable of accurate fibrosis grading. While acknowledging current limitations, ongoing and future research efforts will continue to refine and validate these models, ultimately aiming to improve clinical decision-making and patient outcomes in bone marrow evaluation.

## 4. Resource Availability

### 4.1 Potential Use Cases

#### Diagnostic Assistance

Clinicians can leverage DL models trained on this dataset to assist in the quantitative assessment of fibrosis grade in MPN and other disease conditions. This approach may be particularly valuable in differentiating between MPN, such as the cellular phase of primary myelofibrosis and essential thrombocytosis. Furthermore, it could aid in the early prediction of the risk of MPN patients progressing to the overt myelofibrosis phase. The detection of microfoci of fibrosis, evidenced by increased reticulin deposition, may also facilitate the identification of patchy areas of involvement in patients with lymphoma or metastasis.

#### Prognostic Assessment

The dataset can enhance prognostic evaluations by providing objective measurements of fibrosis severity, which is critical for predicting disease progression and patient outcomes. In patients undergoing a repeat bone marrow biopsy, the dataset can be used to quantitatively reassess fibrosis, thereby predicting responses to therapies/interventions. Furthermore, minimal responses in patients receiving treatments or novel targeted therapies, which might be visually underrepresented, can be documented more precisely. This is particularly crucial in the context of clinical trials, where accurate assessment of therapeutic efficacy is essential.

#### Treatment Planning

Accurate assessment of fibrosis levels can guide treatment decisions, can significantly improve treatment planning by providing objective and quantitative measurements of fibrosis severity, enabling more accurate evaluations of disease progression and patient outcomes. Clinicians can use this data to tailor treatment plans, monitor responses to therapies, and make timely adjustments, particularly in chronic conditions like MPN. Additionally, the ability to detect minimal responses to novel therapies, especially in clinical trials, enhances the assessment of treatment efficacy. By supporting personalized medicine and reducing reliance on subjective interpretations, this approach ensures more consistent and effective patient management.

### 4.2 Licensing

The dataset will be hosted on an open-source platform for access by public. It may be obtained by submission of an online application form and acceptance of a Data Use Agreement. The application must include the investigator’s institutional affiliation and the proposed uses of the BoMBR dataset.

### 4.3 Ethical Considerations

All procedures performed in studies involving human participants were per the ethical standards of the institutional and/or national research committee and with the 1964 Helsinki Declaration and its later amendments or comparable ethical standards. The approval from the Institutional Ethical Clearance Committee, Postgraduate Institute of Medical Education and Research (PGIMER), Chandigarh was obtained for conducting the study.

Subjects’ consent was taken for using their biopsy images for research purposes. Subjects were informed verbally (in their vernacular language) about the option to opt out of the study and only the data for those subjects who opted in was included in the study.

## 5. Methods

After the initial collection and screening of the data, we obtained the digital images for the samples. These samples were then subjected to an automated rough annotation pipeline, followed by manual annotation by trained pathologists. After manual annotation, each of the image contains segmentation masks corresponding to each class viz. cell, fat, bony trabeculae and reticulin fibers.

### 5.1 Data collection

The BMT biopsy data was acquired using standardized procedures at the Postgraduate Institute of Medical Education and Research, Chandigarh (PGIMER). These procedures are as per medical standards as mentioned in Bancroft’s Theory and Practice of Histological Techniques (Suvarna et al., 2012). After aspiration, samples underwent decalcification, fixation and paraffin embedding. Then biopsy specimens were prepared for microscopic examination by sectioning into 2-3 microns samples and staining with reticulin stain based on silver impregnation technique. High-resolution digital images of stained biopsy sections were captured using an Olympus BX53 microscope at a magnification of 400x and 1000x. A total of 19 patients diagnosed with MPN participated in the study. The time taken for data collection varied depending on various factors, including sample preparation and imaging. For sample preparation the time taken was approximately 120 minutes. For taking an image it takes approximately 10 seconds. The annotation of the images depends on the MF grade and ranges from 5 minutes to 20 minutes for MF-0 to MF-3.

In our dataset, we ensured comprehensive representation across all myelofibrosis MF grades, encompassing a diverse range of pathological conditions. Specifically, we included 39 images of MF-0, 47 images of MF-1, 78 images of MF-2, and 37 images of MF-3. Hemorrhages by the leakage of blood from ruptured blood vessels into surrounding tissues or as bone marrow aspiration induced ones, are a common feature observed in bone marrow pathology. Our dataset initially comprised 202 images, but we discarded 1 image due to excessive hemorrhage, leaving us with a total of 201 images. This dataset features 55 images exhibiting hemorrhages, capturing its diverse manifestations within the bone marrow environment. By incorporating a wide range of MF grades and cellular samples, our dataset offers researchers a comprehensive resource for studying the complex interplay between fibrosis, cellular composition, and disease progression in MPN and other disease states. Table 1 provides a comprehensive overview of the BoMBR dataset characteristics

**Table 1:**
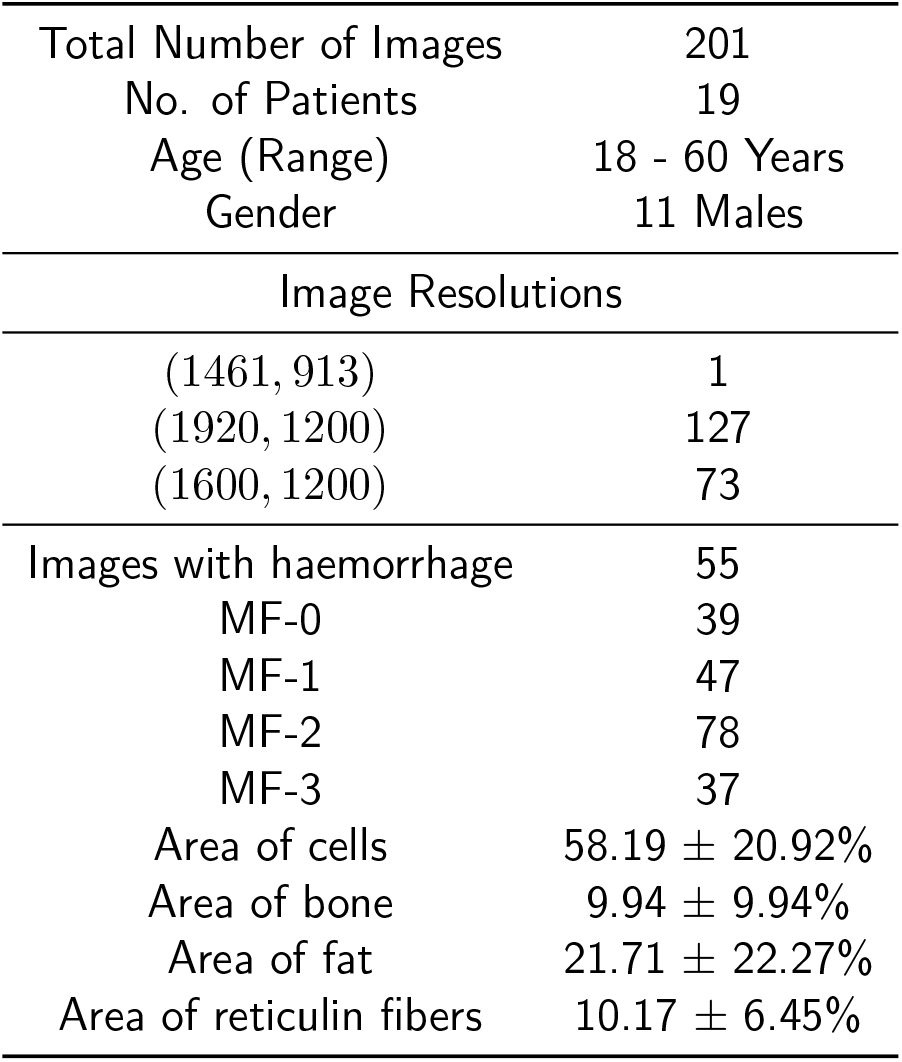
Characteristics of the BoMBR dataset.

|  |  |
| --- | --- |
| Total Number of Images | 201 |
| No. of Patients | 19 |
| Age (Range) | 18 - 60 Years |
| Gender | 11 Males |
| Image Resolutions |  |
| (1461, 913) | 1 |
| (1920, 1200) | 127 |
| (1600, 1200) | 73 |
| Images with haemorrhage | 55 |
| MF-0 | 39 |
| MF-1 | 47 |
| MF-2 | 78 |
| MF-3 | 37 |
| Area of cells | 58.19 ± 20.92% |
| Area of bone | 9.94 ± 9.94% |
| Area of fat | 21.71 ± 22.27% |
| Area of reticulin fibers | 10.17 ± 6.45% |

### 5.2 Data annotation process

To assist in annotating our BMT biopsy specimens, we developed a preprocessing pipeline. We use several image processing techniques to generate a preliminary rough annotation of the image samples. This approach ensures that pathologists do not have to begin annotations from scratch and can focus on refining annotations, including small details such as reticulin fibers. The final annotation is performed manually by the hematopathologist.

#### 5.2.1 Generating instance segmentations in the image

The annotation process for our BMT biopsy specimens began with automated annotation using the Segment Anything Model (SAM) (Kirillov et al., 2023). SAM is a state-of-the-art model designed to generate masks for various components in images, providing initial uncategorized instance segmentation masks for the image. SAM gave us multiple uncategorized masks for all the structures present in the bone marrow samples. These masks were then further refined and categorized based on specific rules tailored to the features of the bone marrow components. Details about the hyperparameters used for the SAM model can be found in section B of the supplementary material.

#### 5.2.2 Bony trabeculae identification

Bony trabeculae are typically visualized in the orange color range in medical images due to their specific staining properties. This characteristic color made it feasible to use image processing approaches for the identification of bony trabeculae.

The process began with the application of color-based segmentation techniques to masks generated by the SAM (Kirillov et al., 2023). The identification process employed two distinct color ranges: orange and black. The orange color range ([0, 50, 20] to [100, 255, 255]; HSV color space), was utilized to capture hues typically associated with bony structures in medical images. Concurrently, a black color range ([0, 0, 0] to [30, 30, 30]) helped in excluding the background or non-bony regions from consideration.

For each mask derived from the segmentation process, the algorithm calculated the proportion of pixels falling within the orange color range relative to the total area (excluding the black pixels). Masks where the proportion of orange pixels exceeded 45% of the non-black area were identified as containing bony trabeculae.

#### 5.2.3 Fat Detection

To identify masks for fat, we began by isolating non-black areas in the masks. This was done using a black color range ([0, 0, 0] to [30, 30, 30], HSV color space). Given that fat regions have consistent color, we then calculated the standard deviation of pixel values in the non-black areas of the masks. Regions where the average standard deviation of pixel intensities value was less than or equal to 20 were identified as fat due to their consistent color characteristics.

#### 5.2.4 Cell Segmentation

The remaining masks, excluding bony-trabeculae and fat, were categorized as cell regions. This broad category encompassed various cellular structures (including reticulin) in the bone marrow samples. Once the different masks were assigned categories, we aggregated all the masks by superimposing them on top of each other, thereby obtaining a single mask for each category.

#### 5.2.5 Reticulin Identification

The process of identifying reticulin fibers involved manual thresholding of the combined cell mask using OpenCV’s cv2.threshold function (Bradski, 2000). Due to varying illumination levels in the images, the threshold values were varied manually (150 to 180) in order to maximize reticulin identification. However, since a single constant threshold value was insufficient to comprehensively identify reticulin, we applied a two-step process to minimize false positives. We utilized the fact that reticulin fibers are elongated, fiber-like structures, whereas nuclei are circular and have low elongation values close to 0. We, thus, used the elongation factor to distinguish between reticulin and nuclei using the following steps:

- Bounding Box Generation: Bounding boxes were generated around the identified components. The elongation of each bounding box was calculated using the following formula:

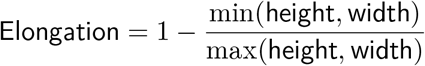 This formula quantified the degree of elongation based on the dimensions of the bounding box. If either the width or height of the bounding box was zero, indicating a non-elongated structure, the elongation was set to zero.
- Contour Filtering: Contours with elongation values exceeding a threshold of 0.25 were categorized as reticulin. This step ensured that only elongated structures, characteristic of reticulin fibers, were retained while excluding non-relevant components like nuclei, which are typically circular.

After generating the reticulin mask, it was subtracted from the original cell mask to produce a cell mask without reticulin. The final set of annotations included masks for reticulin, fat, bony trabeculae, and cells without reticulin.

### 5.3 Annotation by Experts

Despite the advantages of automated annotation, the rough annotations could not be considered precise due to certain inaccuracies. For instance, the SAM encountered challenges in generating precise masks for fat regions having small area, due to which these occasionally got included in the cell mask. Moreover, despite efforts to exclude them using elongation checks, nuclei and other cellular structures close to reticulin fibers were included in the same bounding box as reticulin fibers. To address these issues and ensure the quality of annotations, manual verification was essential. The manual annotation process involved two hematopathologists who were trained on our software for performing manual annotation. Experts reviewed each image, refining the annotations and correcting any discrepancies identified during the automated phase. Each image was annotated by the hematopathologist taking approximately 5 to 20 minutes for its complete annotation. Examples of images along with their expert annotations can be found in section C of the supplementary material.

To ensure consistency and compatibility with other tools and platforms, all annotations were converted into Pascal VOC format. This standardized format facilitated seamless integration with existing frameworks and enhanced the accessibility of the annotated dataset for further research and development. CVAT (Sekachev et al., 2020) was chosen for its intuitive user interface and open-source nature, facilitated this manual annotation process.

In summary, the annotation process for the bone marrow biopsy images combined automated methods with manual verification to achieve accurate and reliable annotations. This hybrid approach leveraged the strengths of automated segmentation techniques while mitigating potential inaccuracies through expert intervention. The resulting annotated dataset, available in Pascal VOC format and annotated using CVAT, serves as a valuable resource for research and development in bone marrow evaluation and related fields.

## 6. Using BoMBR Dataset for Marrow Fibrosis Grade Prediction

### 6.1 Motivation

MPN are a group of disorders characterized by acquired mutations in hematopoietic stem cells, affecting the MPL-JAK-STAT (Myeloproliferative Leukemia Virus-Janus Kinase-Signal Transducer and Activator of Transcription) signaling pathway and leading to excessive proliferation of one or more blood cell lineages (Sabattini et al., 2021; Gianelli et al., 2017; Rampal et al., 2014; O’Sullivan and Mead, 2019). Despite advances in understanding MPN, the precise mechanisms underlying the initiation and progression of MF remain poorly understood. The significance of evaluating fibrosis levels in MPN is deeply integrated into the classification system established by the World Health Organization (WHO) for common MPN, including essential thrombocythemia (ET), polycythemia vera (PV), primary myelofibrosis (PMF), and pre-fibrotic primary myelofibrosis (pre-PMF) (Arber et al., 2016). Fibrosis severity plays a pivotal role in the clinical management of MPN, with minor fibrosis (MF-1) observed in PV linked to poorer survival outcomes, while more advanced fibrosis is associated with a complex karyotype (Barbui et al., 2012; Boiocchi et al., 2013). Escalating levels of fibrosis correspond to worsening hematological and clinical parameters, as well as overall prognosis (Gianelli et al., 2012; Vener et al., 2008; Thiele and Kvasnicka, 2006). The current method for assessing bone MF relies heavily on manual grading by pathologists.

### 6.2 Dataset Preparation

The dataset was split into training and validation sets using an 80:20 ratio, stratified across all MF grades to ensure balanced representation giving us a split of 161 train images and 40 validation images. For the test set, we captured an additional 50 images from 10 additional subjects (other than the train set) and annotated them to prevent data leakage among subjects or from the initial dataset to the test set.

### 6.3 Model Used

For the segmentation task, we used a UNet (Ronneberger et al., 2015) model with an Xception (Chollet, 2017) backbone. The encoder we used was taken from Keras (Chollet et al., 2015) and was pretrained on ImageNet (Deng et al., 2009) weights with an input size of 512 × 512. The model architecture included residual blocks and skip connections, totaling 12 residual blocks and 4 skip connections. The architecture of the decoder was adopted from Kaggle. Dice loss was used as the loss function, while monitoring the Dice coefficient and mean IoU. We applied dropout for regularization to enhance feature learning. We used a learning rate of 10^−4^ with an AdamW optimizer, a batch size of 8, and set the maximum number of epochs to 200.

For the classification task, we used a CNN model with an encoder based on the Xception backbone. We used a learning rate of 10^−5^ with an Adam optimizer, a batch size of 8, and set the maximum number of epochs to 70. The Xception model layers were frozen to retain pretrained weights, and additional convolutional and dense layers were incorporated for classification.

For both models, we implemented early stopping with a patience of 10 using the TensorFlow Python library (TensorFlow, 2018), running on a NVIDIA T1000 8GB GPU. Additionally, we utilized a learning rate scheduler with a reduction factor of 0.8, monitoring accuracy, and allowing a patience of 4 epochs before reducing the learning rate to a minimum of 10^−8^.

It is to be noted that we performed these simulations to show that robust models can be built on top of our datasets performing meaningful tasks that lead to the reduction of clinical efforts and more accurate performance. We, therefore, focused on building suitable model architectures instead of optimizing model performances. Further research and models can be explored to optimize the model performance.

### 6.4 Results

#### 6.4.1 Segmentation

We conducted multi-class segmentation using a Xception model pretrained on the ImageNet dataset, where we froze the encoder weights and trained the decoder from scratch until the training loss converged (Figure 4). Details of the model’s performance are summarized in Table 2. A sample segmentation result is illustrated in Figure 5, showcasing the model’s ability to accurately classify pixels into 4 distinct classes. The segmentation model demonstrated robust performance, achieving a Dice score of 0.9230, indicating its capability to discern meaningful patterns within the dataset. Additionally, graphical representations of the training and testing Dice Coefficient and mIoU over 200 epochs are provided in section D of the supplementary material.

**Table 2:**
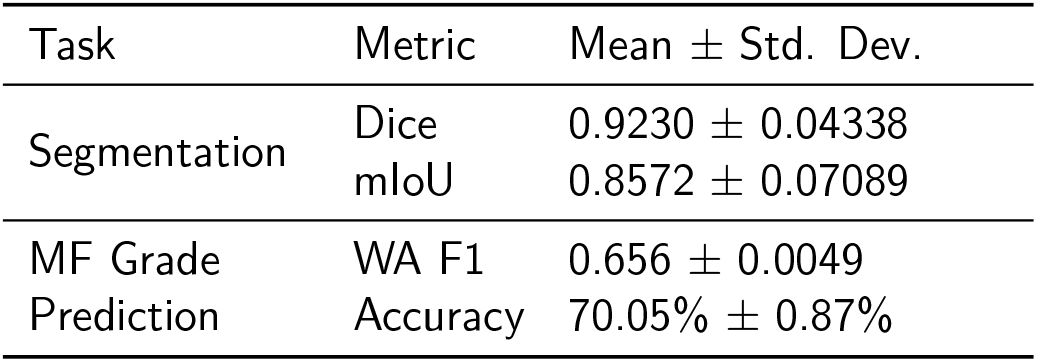
Description of model performance across five different random initializations. Abbreviations used: Weighted Average (WA).

**Figure 4.**
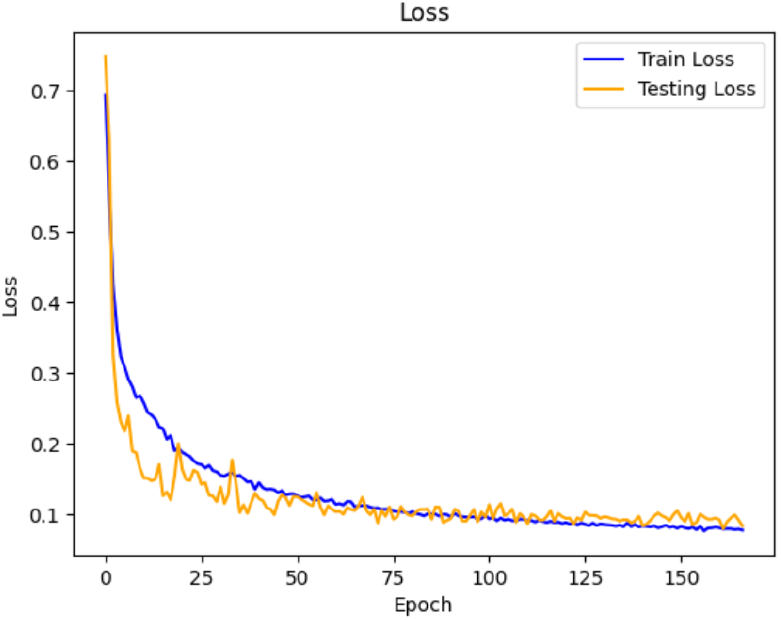
Plot of the dice loss over the number of epochs for the train and test sets for the segmentation model.

**Figure 5.**
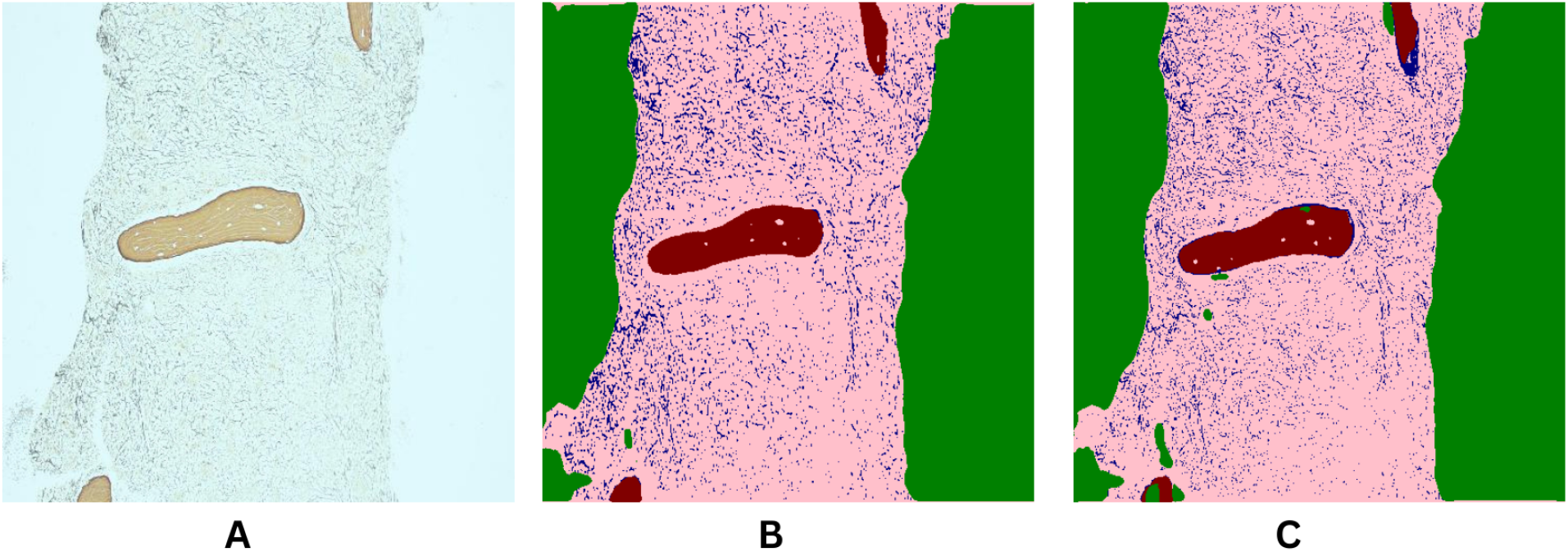
Segmentation model’s labels for a test sample. A: Sample image from the test set; B: Image with ground truth masks overlaid on top; and C: Image with predicted segmentation masks from our trained model. Note that color pink represents cell, red represents bone, blue represents reticulin, and green represents fat.

#### 6.4.2 Fibrosis Grade Prediction

For fibrosis grade prediction, we propose a convolutional neural network (CNN) architecture. The network utilizes a pre-trained Xception encoder on ImageNet for initial feature extraction. It takes masked grayscale images of size 512*×* 512, where only reticulin is present and the background is blacked out. These images are derived from reticulin masks predicted using our segmentation model. These extracted features undergo processing through an additional convolutional layer with 512 filters and a 3*×* 3 kernel size to refine spatial information. Subsequently, global average pooling aggregates these refined features, which are then passed through a dense layer comprising 128 units with ReLU activation. A dropout layer is applied for regularization before a final dense layer with softmax activation predicts the probabilities for the four myelofibrosis (MF) grades. Performance metrics for this classification task are detailed in Table 2. We found that we were able to predict the MF grade with a mean accuracy of 87.5% (over 5 folds). The section E of the supplementary material contains the detailed classification report and confusion matrix for this model.

It is to be noted that although the ground truth MF grades were based on the consensus of two hematopathologists, they are subjective and often cases that are borderline. Further investigation based on uncertainty quantification methods can be performed to improve our results.

## Acknowledgments

We acknowledge important conversations with our colleagues Amit Kumar and Maninder Kaur.

## Ethical Standards

The work follows appropriate ethical standards in conducting research and writing the manuscript, following all applicable laws and regulations regarding treatment of animals or human subjects.

## Data availability

The dataset can be found on an open-source platform for public access. We followed the rules and regulations as mentioned in Starmans and Tsirikoglou (2024) and uploaded the dataset on Raina et al. (2024). To obtain the dataset, users must submit an online application form and accept the Data Use Agreement. The application must include the investigator’s institutional affiliation and the proposed uses of the BoMBR dataset.

