## Supplementary Materials for "BoMBR: An Annotated Bone Marrow Biopsy Dataset for Segmentation of Reticulin Fibers"

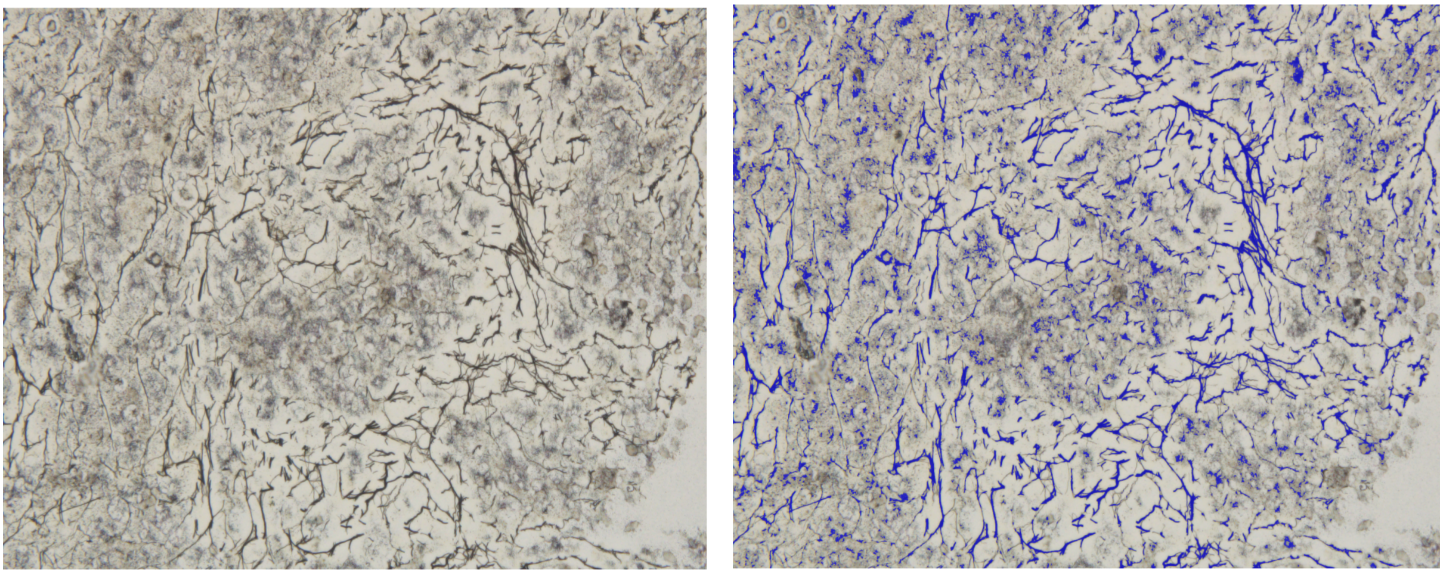

A B

Figure 1: Sample of a BMT image from our dataset. A: Original unsegmented image B: Annotated image with reticulin fibers labeled in blue color

accurate segmentation and quantification difficult (Lucero et al., 2016). For example, Teman et al. (2010) developed a computer-aided solution that utilizes color deconvolution techniques to isolate the reticulin fibers from the image and then manually grade the MF. This work, however, ignores the complex polychromatic nature of reticulin fibers and is not fully automated thereby having a manual grading component.Lucero et al. (2016) quantified bone MF in a mouse model of myelofibrosis. However, its applicability to human bone marrow biopsies and more complex disease presentations requires further validation and adaptation. More recently, Ryou et al. (2023) introduced Continuous Indexing of Fibrosis (CIF), a DL approach aimed at enhancing the quantification and monitoring of fibrosis in using bone marrow samples. CIF utilizes a continuous 0-1 scale, departing from the traditional discrete 0-3 grading system, thereby improving fibrosis assessment in Myeloproliferative Neoplasms (MPN). However, a potential drawback is that this new grading scale may not align with established clinical practices, posing challenges for clinicians in interpreting and applying the results in standard clinical settings. As a result, these computer-aided approaches may not provide reliable and reproducible results, further highlighting the need for alternative methods that can overcome these limitations and offer more robust assessments of bone MF.

| | |
|---|---|
| Total Number of Images | 201 |
| No. of Patients | 19 |
| Age (Range) | 18 - 60 Years |
| Gender | 11 Males |
| Image Resolutions | |
| $(1461, 913)$ | 1 |
| $(1920, 1200)$ | 127 |
| $(1600, 1200)$ | 73 |
| Images with haemorrhage | 55 |
| MF-0 | 39 |
| MF-1 | 47 |
| MF-2 | 78 |
| MF-3 | 37 |
| Area of cells | 58.19 $\pm$ 20.92% |
| Area of bone | 9.94 $\pm$ 9.94% |
| Area of fat | 21.71 $\pm$ 22.27% |
| Area of reticulin fibers | 10.17 $\pm$ 6.45% |

Table 1: Characteristics of the BoMBR dataset

- Bounding Box Generation: Bounding boxes were generated around the identified components. The elongation of each bounding box was calculated using the following formula:

  $$\text{Elongation} = 1 - \frac{\min(\text{height}, \text{width})}{\max(\text{height}, \text{width})}$$

  This formula quantified the degree of elongation based on the dimensions of the bounding box. If either the width or height of the bounding box was zero, indicating a non-elongated structure, the elongation was set to zero.

- Contour Filtering: Contours with elongation values exceeding a threshold of 0.25 were categorized as reticulin. This step ensured that only elongated structures, characteristic of reticulin fibers, were retained while excluding non-relevant components like nuclei, which are typically circular.

After generating the reticulin mask, it was subtracted from the original cell mask to produce a cell mask without reticulin. The final set of annotations included masks for reticulin, fat, bony trabeculae, and cells without reticulin.

#### 5.3 Annotation by Experts

Despite the advantages of automated annotation, the rough annotations could not be considered precise due to certain inaccuracies. For instance, the SAM encountered challenges in generating precise masks for fat regions having small area, due to which these occasionally got included in the cell mask. Moreover, despite efforts to exclude them using elongation checks, nuclei and other cellular structures close to reticulin fibers were included in the same

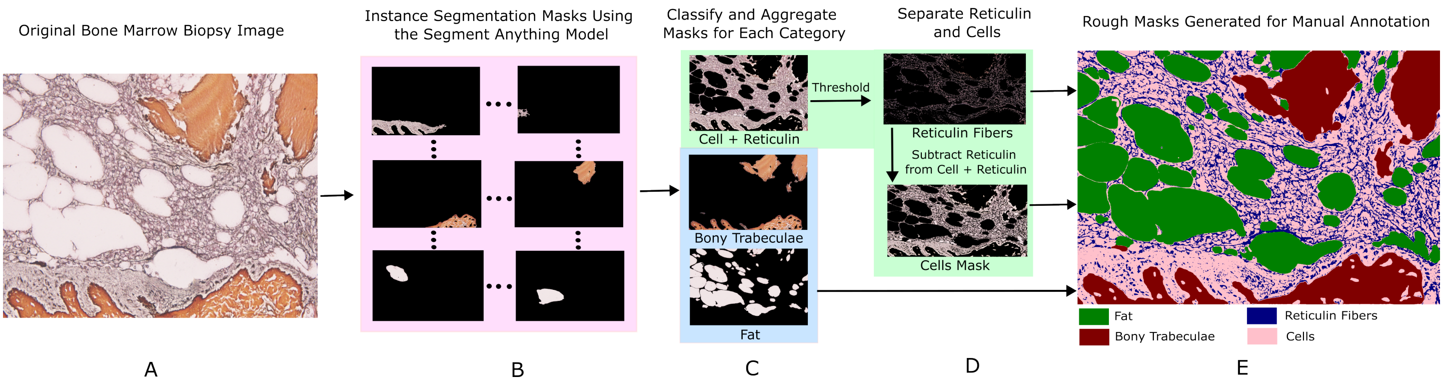


Figure 2: Flowchart representation of automatic annotations done by SAM. A: Original Image. B: Generating masks and classifying them: Masks with pixel intensity in the orange color range are classified as bone masks, masks having consistency in color among the remaining masks are considered as fat masks, and the remaining masks are categorized as cell masks. C: Masks of every category are added together to form one mask per category. D: Performing thresholding and shape filtering to isolate reticulin, resulting in a reticulin mask and a cell mask without reticulin. E: Annotations generated using SAM and classification.

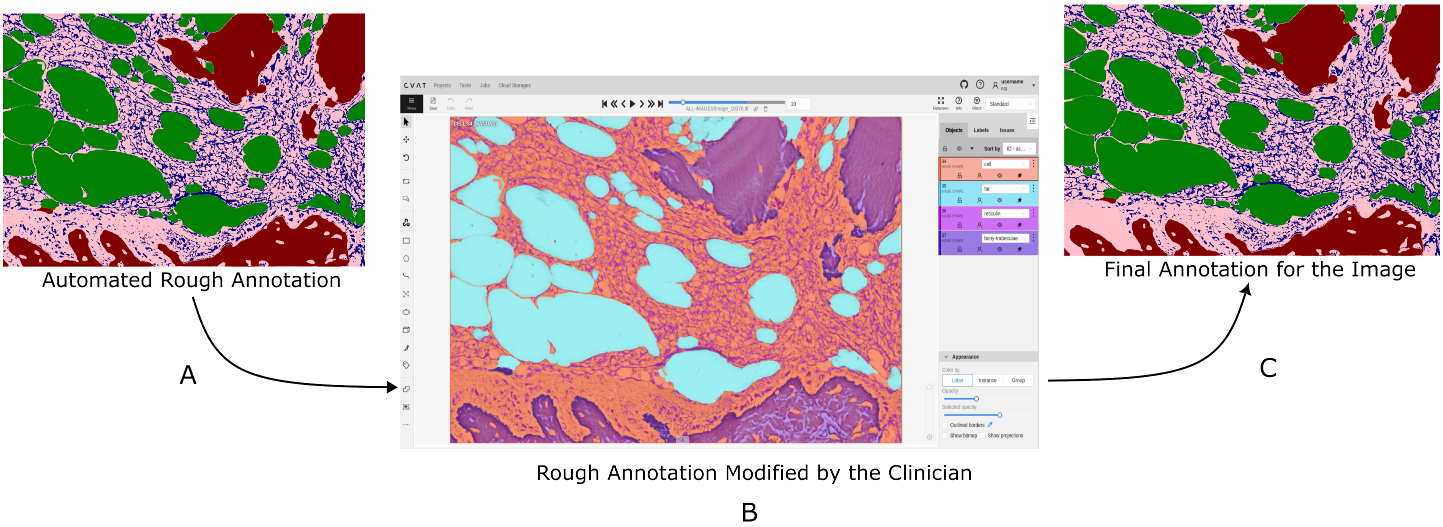


Figure 3: Procedure for annotation by experts. A: Automatic annotation using SAM and classification of masks. B: The pathologist uses Computer Vision Annotation Tool (CVAT) for refining annotations. C: Final annotation

bounding box as reticulin fibers.

#### 6.4 Results

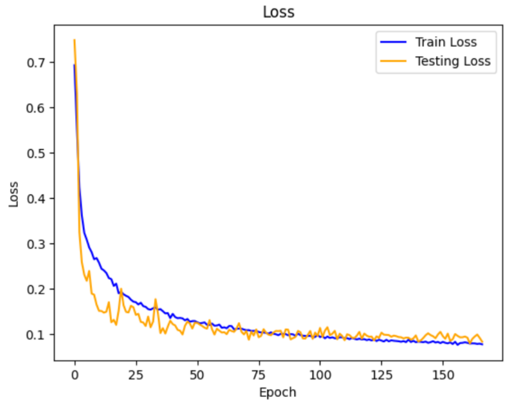


Figure 4: Plot of the dice loss over the number of epochs for the train and test sets for the segmentation model.

##### *6.4.1 Segmentation*

We conducted multi-class segmentation using a Xception model pretrained on the ImageNet dataset, where we froze the encoder weights and trained the decoder from scratch until the training loss converged (Figure 4). Details of the model's performance are summarized in Table 2. A sample segmentation result is illustrated in Figure 5, showcasing the model's ability to accurately classify pixels into 4 distinct classes. The segmentation model demonstrated robust performance, achieving a Dice score of 0.9230, indicating its capability to discern meaningful patterns within

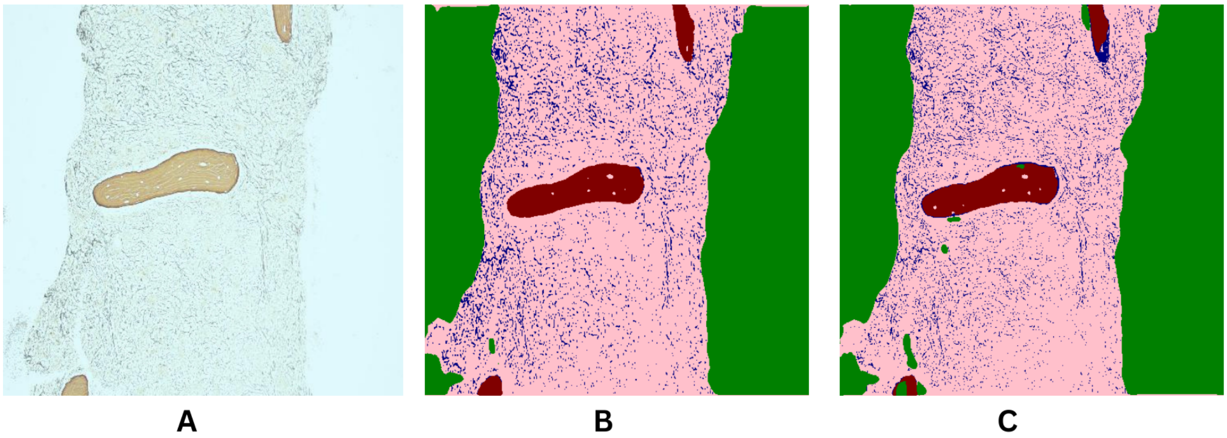


Figure 5: Segmentation model's labels for a test sample. A: Sample image from the test set; B: Image with ground truth masks overlaid on top; and C: Image with predicted segmentation masks from our trained model. Note that color pink represents cell, red represents bone, blue represents reticulin, and green represents fat.

the dataset. Additionally, graphical representations of the training and testing Dice Coefficient and mIoU over 200 epochs are provided in section D of the supplementary material.

#### *6.4.2 Fibrosis Grade Prediction*

For fibrosis grade prediction, we propose a convolutional neural network (CNN) architecture. The network utilizes a pre-trained Xception encoder on ImageNet for initial feature extraction. It takes masked grayscale images of size $512 \times 512$, where only reticulin is present and the background is blacked out. These images are derived from reticulin masks predicted using our segmentation model. These extracted features undergo processing through an additional convolutional layer with 512 filters and a $3 \times 3$ kernel size to refine spatial information. Subsequently, global average pooling aggregates these refined features, which are then passed through a dense layer comprising 128 units with ReLU activation. A dropout layer is applied for regularization before a final dense layer with softmax activation predicts the probabilities for the four myelofibrosis (MF) grades. Performance metrics for this classification task are detailed in Table 2. We found that we were able to predict the MF grade with a mean accuracy of 87.5% (over 5 folds). The section E of the supplementary material contains the detailed classification report and confusion matrix for this model.

It is to be noted that although the ground truth MF grades were based on the consensus of two hematopathologists, they are subjective and often cases that are borderline. Further investigation based on uncertainty quantification methods can be performed to improve our results.

| Task | Metric | Mean $\pm$ Std. Dev. |
|---|---|---|
| Segmentation | Dice | $0.9230 \pm 0.04338$ |
| | mIoU | $0.8572 \pm 0.07089$ |
| MF Grade Prediction | WA F1 | $0.656 \pm 0.0049$ |
| | Accuracy | $70.05\% \pm 0.87\%$ |

Table 2: Description of model performance across five different random initializations. Abbreviations used: Weighted Average (WA).

### Acknowledgments

We acknowledge important conversations with our colleagues Amit Kumar and Maninder Kaur.

### Ethical Standards

The work follows appropriate ethical standards in conducting research and writing the manuscript, following all applicable laws and regulations regarding treatment of animals or human subjects.

### Conflicts of Interest

The authors declare that they have no conflicts of interest.

### Data availability

The dataset can be found on an open-source platform for public access. We followed the rules and regulations as mentioned in Starmans and Tsirikoglou (2024) and uploaded the dataset on Raina et al. (2024). To obtain the dataset, users must submit an online application form and accept the Data Use Agreement. The application must include the investigator's institutional affiliation and the proposed uses of the BoMBR dataset.

Developers TensorFlow. Tensorflow. *Site oficial*, 2018.

J Thiele and HM Kvasnicka. Grade of bone marrow fibrosis is associated with relevant hematological findings—a clinicopathological study on 865 patients with chronic idiopathic myelofibrosis. *Annals of hematology*, 85:226–232, 2006.

Claudia Vener, Nicola Stefano Fracchiolla, Umberto Gianelli, Rossella Calori, Franca Radaelli, Alessandra Iurlo, Sabrina Caberlon, Giancarla Gerli, Leonardo Boiocchi, and Giorgio Lambertenghi Deliliers. Prognostic implications of the european consensus for grading of bone marrow fibrosis in chronic idiopathic myelofibrosis. *Blood, The Journal of the American Society of Hematology*, 111 (4):1862–1865, 2008.
